# Detection of Locomotion Deficit in a Post-Traumatic Syringomyelia Rat Model Using Automated Gait Analysis Technique

**DOI:** 10.1101/2021.05.19.444781

**Authors:** Dipak D. Pukale, Mahmoud Farrag, Nic D. Leipzig

## Abstract

Syringomyelia (SM) is a spinal cord disorder in which a cyst (syrinx) filled with fluid forms in the spinal cord post-injury/disease, in patients syrinx symptoms include loss of pain and temperature sensation or locomotion deficit. Currently, there are no small animal models and connected tools to help study the functional impacts of SM. The objective of this study was to determine the detectability of subtle locomotion deficits due to syrinx formation/expansion in post-traumatic syringomyelia (PTSM) rat model using the recently reported method of Gait Analysis Instrumentation, and Technology Optimized for Rodents (GAITOR) with Automated Gait Analysis Through Hues and Areas (AGATHA) technique. First videos of the rats were collected while walking in an arena (using GAITOR) followed by extracting meaningful locomotion information from collected videos (using AGATHA protocol. PTSM injured rats demonstrated detectable locomotion deficits in terms of duty factor imbalance, paw placement accuracy, step contact width, stride length, and phase dispersion parameters compared to uninjured rats due to SM. We concluded that this technique could detect mild and subtle locomotion deficits associated with PTSM injury, which also in future work could be used further to monitor locomotion responses after different treatment strategies for SM.

## 1 Introduction

Syringomyelia (SM) is a neurological disorder where a cerebrospinal fluid (CSF)-filled cyst, also known as a syrinx, forms in the spinal cord due to multiple acquired and congenital causes. It is a common coincidence with multiple neurological diseases such as Chiari malformation I, trauma, and several other disorders (1,2). SM associated with spinal cord injury (SCI) is called post-traumatic SM (PTSM), and it develops in about 21-28% of people with SCI (1). It is not always easy to track the triggering trauma, as it sometimes results from minor injury and takes months to years to fully develop (3), making it challenging to prevent. SM exhibits mainly with chronic progressive pain and neurological deficits (1) because syrinxes compress the surrounding neural tissue while they expand and enlarge, undesirably impacting patient quality of life. Current treatment options for SM are mainly surgical interventions such as shunting, duraplasty, adhesiolysis, or decompression (4–6). Even though surgical treatments are standard options to treat SM, due to surgical complications and their high failure rate (>80%) (6), these options do not provide enough confidence to clinicians to treat SM acceptably. Multiple competing theoretical mechanisms have attempted to explain syrinx formation/expansion (7–11). More recent work, including our own, has explored molecular aspects of SM to attempt to better understand the connections to SM pathology (12–16) while also extending this to animal PTSM models (10,15,17).

Symptoms of SM include a distorted sense of pain and temperature and progressive weakness in the back, shoulders, arms, or legs, along with bowel and bladder dysfunction, which correlate to the presence of dynamic syrinxes in the spinal cord. These symptoms depend, in large part, on the size and position of syrinxes in the spinal cord (3,18,19). To facilitate SM research towards developing non-surgical treatment options it is pivotal to establish functional techniques to monitor the sensory or motor activities concerning those potential treatments in the future. There have been limited reported attempts that have evaluated locomotion deficits in SM animals, with one recent report showing that pet Cavalier King Charles Spaniels with Chiari 1 and SM have an increased variation of the ipsilateral distance between paws, varied length of the stride and a wider base of support in the hind limbs (20). Perhaps because PTSM generated in rodent models is an indirect and non-severe injury, no gait analysis technique thus far has been able to detect locomotion deficits. Recently, we reported a new method called Gait Analysis Instrumentation, and Technology Optimized for Rodents (GAITOR) with Automated Gait Analysis Through Hues and Areas (AGATHA) (21). The AGATHA with GAITOR technique has previously been used to determine locomotion changes in several injury models, including a sciatic nerve injury model, an elbow joint contracture model, SCI models, and an osteoarthritis joint pain model (21–24). The established ability to obtain and tune SCI-specific locomotion parameters motivated us to apply this technique for a PTSM rat model.

The purpose of this study was to detect and ultimately determine locomotion deficits that coincide with syrinx formation/expansion/elongation in the spinal cord using a PTSM rat model by applying GAITOR with AGATHA. Since PTSM or other SM animal models are non-severe and indirect there is no agreed-upon technique able to reliably detect functional outcomes in terms of motor and/or sensory functions. The findings from this study will be useful to provide needed tools to better facilitate SM research toward better treatments and diagnoses in the near future.

## 2 Materials and Methods

### 2.1 PTSM Injury model

All experimental manipulations related to animals were conducted according to the University of Akron Institutional Animal Care and Use Committee. Two groups were utilized in the study: (1) healthy rats with no surgical procedures received (uninjured group) and (2) animals with post-traumatic syringomyelia (PTSM group). 10 week old male Wistar rats were used for this study, including 6 rats in the uninjured group and 9 rats in PTSM groups. PTSM induction surgery involved an intraspinal injection of 2 μL of 24 mg/mL quisqualic acid (QA) (Enzo Life Sciences, Farmingdale, NY) and 5μL of 250 mg/mL kaolin (Avantor, Center Valley, PA, USA) in the subarachnoid space after laminectomy for animals in the PTSM group. While QA injection-induced excitotoxicity, kaolin injections were used to create subarachnoid adhesion and increased CSF pressure. The two events synergistically lead to syrinx formation in the spinal cord. These injections were made in the right dorsal quadrant of the cord between levels C7 and C8. The combination of local neuron destruction using QA and arachnoiditis caused due to subarachnoid space blockage using kaolin forms a fluid-filled cavity that enlarges over 6 weeks (14,15,17,25). After PTSM injury, functional observations were recorded until the 6-week endpoint, when animals were perfused with PBS followed by a 4% paraformaldehyde (PFA) in PBS solution to displace blood from the tissue and to fix the tissue, respectively.

### 2.2 Immunohistochemistry

Spinal cords were immediately dissected and placed in 4% PFA for 4 h at 4°C for post-fixation, then transferred in 0.2 M phosphate buffer and left overnight. Next, spinal cords were stored in 30% sucrose for 3 days at 4°C before embedding those into the OCT compound (Tissue-Tek). Embedded blocks were stored at −80°C until cryosectioning using a cryostat (Leica CM 1850). 25 μm transverse sections were made for immunohistochemistry and syrinx confirmation. The monoclonal GFAP antibody (1:200, Santa Cruz Biotechnology, Dallas, TX, USA) was used to identify astrocyte cells around the syrinx. Cell nuclei were stained with 10 mM Hoechst 33342 for 7 min followed by mounting using ProLong™ Gold Antifade (ThermoFisher, Waltham, MA, USA). Images were taken using an Olympus FV 1000 confocal microscope.

### 2.3 Automated gait analysis

The detailed description and functions of instrumentation and MATLAB-based AGATHA software have been explained previously (21), and detailed SCI-specific locomotion parameters were discussed in references (21–23). In short, the GAITOR setup (Figure 1) is comprised of two parts, an arena where rats walk back and forth unprompted while videos are recorded using a high-speed camera. A 45° inclined mirror to capture the ventral view of the rat walking on the arena is also included. Next, the files are processed with AGATHA to first filter the background of each rat in the sagittal view and the paws in the ventral view. AGATHA provides the capability to exclude the regions where the rat touches the arena surface other than paws such as with its nose or tail. The program also determines the foot-strike toe-off (FSTO) diagram based on the earliest (foot-strike) and last (toe-off) frames in which a paw is in contact with the arena surface as shown in Figure 1f. Based on the information processed, AGATHA calculates temporal parameters such as duty factor imbalance, phase dispersion, and spatial parameters such as paw placement accuracy, step width, stride length, which provide locomotive information of the animals walking on the arena as we have previously conducted for a hemisection SCI rat model (22,23).

**Figure 1.**
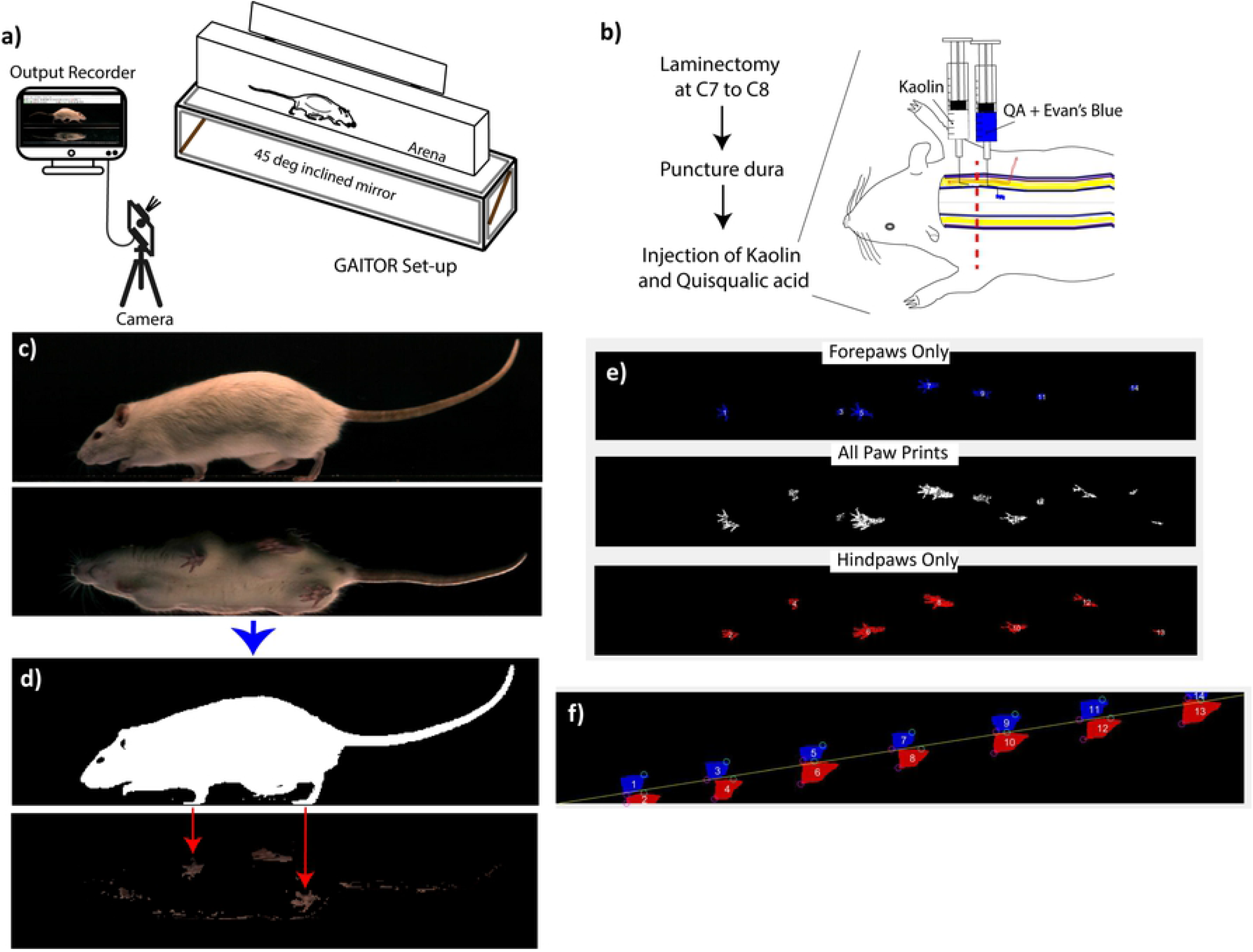
GAITOR set up, injury model, and overview of GAITOR analysis. a) Schematic representation of GAITOR reproduced with permission from the publisher from (22). b) Schematic representation of the PTSM injury at the C7-C8 level of the spinal cord, which potentially impacts forelimb as well as hindlimb movement, to some extent, by forming syrinxes in the spinal cord. c) A representative frame of a GAITOR video showing a sagittal and ventral view of the rat. d) AGATHA software filters rat and paws from the background in the sagittal and ventral view, respectively. e) Pawprint images of forepaws, hind paws, and combined as generated for each video. f) Plots of forepaws (blue) and hind paws (red) showing foot stride (green circle) and toe-off (pink circle) generated for every video.

### 2.4 Thermal allodynia testing

Quantitative measurement of rat responses to thermal stimuli is a known technique in functional assessment in neurological evaluation. A Plantar Analgesia Meter (IITC Inc., Life Science, CA, USA) enables quantification of nociception, which is built based upon the modified Hargreaves method. To perform the test, rats were acclimated in acrylic animal enclosures for 30 min on three consecutive days before testing. Also, every testing session was proceeded by an acclimation period of 5-10 min. Testing was performed the week before and a week after the surgery. The temperature and humidity of the testing room were always checked to ensure consistency in the testing conditions. Testing was done by exposing the center of the plantar surface of the paws to a focused radiant heat light source until the rat retracts its paw or 20 s pass. To time the latency interval, an electronic timer was employed to match the testing light, so both turn on and off simultaneously. A light intensity of 34% was used with a cutoff exposure time of 20 s when it automatically turns off. This corresponds to a glass temperature between 35 – 50°C. Paw withdrawal response was considered when retraction of the paw was accompanied by paw shaking, paw licking, startle, or vocalization. One paw was tested in all rats, then after testing all rats, another paw was tested to make sure that rats were not exposed to unnecessary discomfort that may lead to a stress-induced response.

### 2.5 Experimental design and data processing/stat analysis

Our primary hypothesis was that GAITOR with AGATHA technique would detect mild and subtle locomotion deficit related to syrinx presence in the PTSM injured rat spinal cord. The gait parameter measurements extracted using AGATHA from collected gait videos over time were tested using two-factor ANOVA with Tukey’s *post hoc* (*p*<0.05) for all gait parameters using Minitab software. Uninjured and PTSM groups were compared over 6 week time points to understand the locomotion deficit every week concerning syrinx formation and expansion. The sample size of 6 for uninjured and 9 for PTSM groups were used for the collection of the observations. The gait observations have been collected every week (except week 5). The velocity dependence of the parameters was evaluated by plotting the uninjured rats’ collective data (including all timepoints) for respective parameters vs. velocity. Those parameters which are velocity-dependent are presented in terms of velocity-weighted residuals at each timepoint.

## 3 Results

To begin with, syrinxes in all PTSM rats were confirmed by IHC in injured spinal cord sections and compared with uninjured sections (Figure 2). Using fixed and sectioned tissues a syrinx is characterized by an open cell-free space with a border of dense GFAP positive astrocytes. Confirming syrinx presence in the injured spinal cord is important since our main objective was to determine if locomotion deficits are detectable as a consequence of PTSM injury. Syrinxes fully develop after week 6 in the PTSM injured spinal cord as shown in previous studies including our own (15,17). Using the same animal injury model we recently evaluated syrinx size using a micro CT technique and reported syrinx total surface area in the range of 20 – 40 mm^2^ and total syrinx volume in the range of 1 – 4 mm^3^ (25).

**Figure 2.**
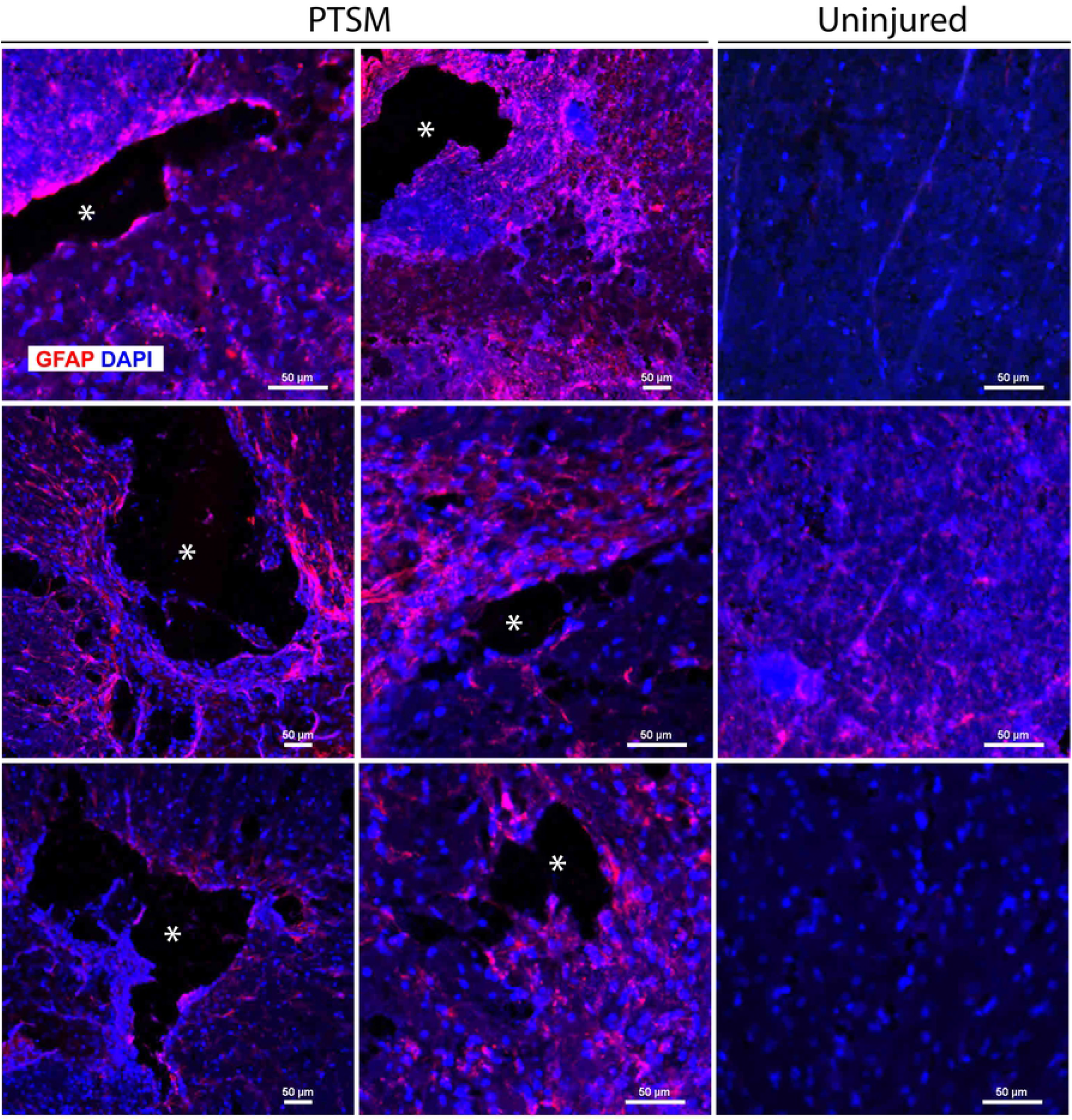
IHC confirmation of cyst/syrinx formation in the spinal cord. Cyst formation within the spinal cord from PTSM group as indicated by GFAP positive cells (astrocyte marker) surrounding the cyst and compared with the uninjured group. * represents cyst

### 3.1 Modified Hargreaves test

Before applying the detailed locomotion deficit technique (AGATHA with GAITOR), we attempted to detect sensory changes caused due to PTSM injury using a modified Hargreaves method as shown in Figure 3. As previously mentioned, the ability to sense temperature and pain in arms and legs coincides with the progression of SM in human patients. The observations recorded in this study using this method showed that there was no statistical difference between PTSM and uninjured groups for all four limbs. This indicates that either PTSM injury does not hamper sensory functions or the modified Hargreaves test is not sensitive enough to capture lost sensory function in a less severe SCI rat model like PTSM.

**Figure 3:**
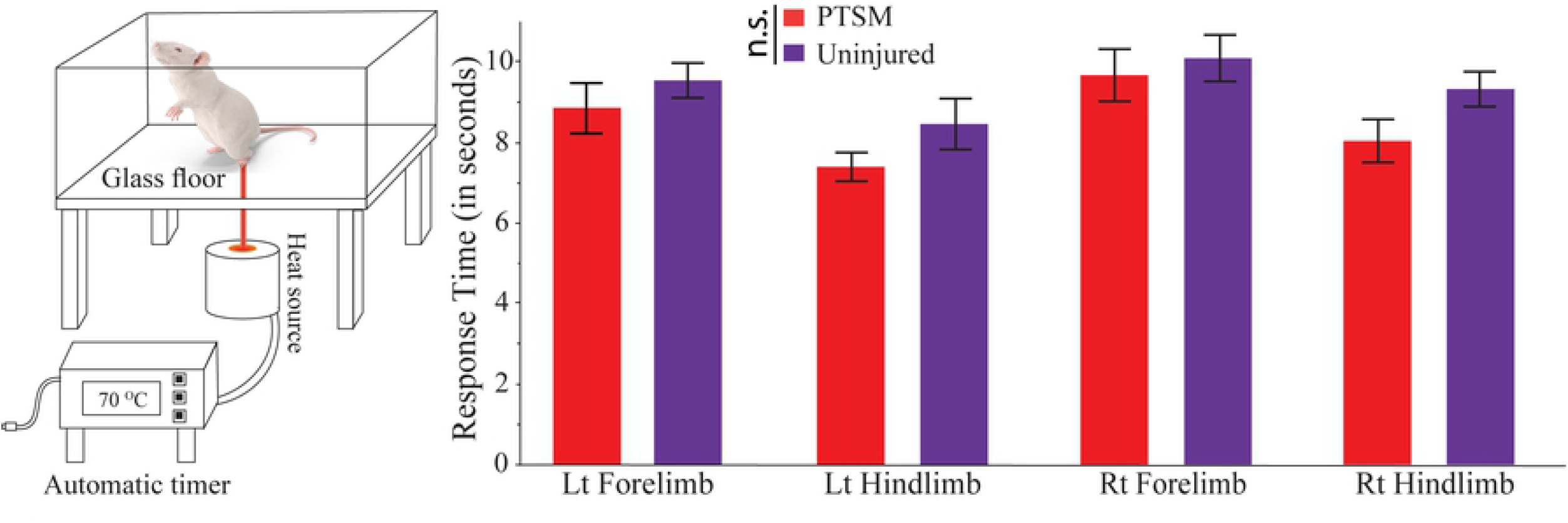
Modified Hargreaves method to test for thermal allodynia. A comparison between uninjured and PTSM rats. The time between animal exposure to thermal stimuli and their animal response was not statistically different between the two groups. The measurements were done in each of the four limbs where the *p*-value for left forelimb is 0.379, for left hindlimb is 0.146, for right forelimb is 0.626, and for right hindlimb is 0.067. Data are presented as mean ± standard error of mean and statistics were determined pairwise via Student’s t-test.

### 3.2 Locomotion parameters

Based on velocity dependency analysis (plotting gait parameters data of uninjured rats at all timepoints vs. velocity), we identified that duty factor, phase distribution are temporal gait parameters. In contrast, paw placement accuracy, step width, and stride length are spatial gait parameters. The velocity-dependent gait parameters (i.e., spatial parameters) are presented in terms of velocity-weighted residuals at each time point (Figures 4–6) similar to the results we have presented in our previous study (23).

**Figure 4.**
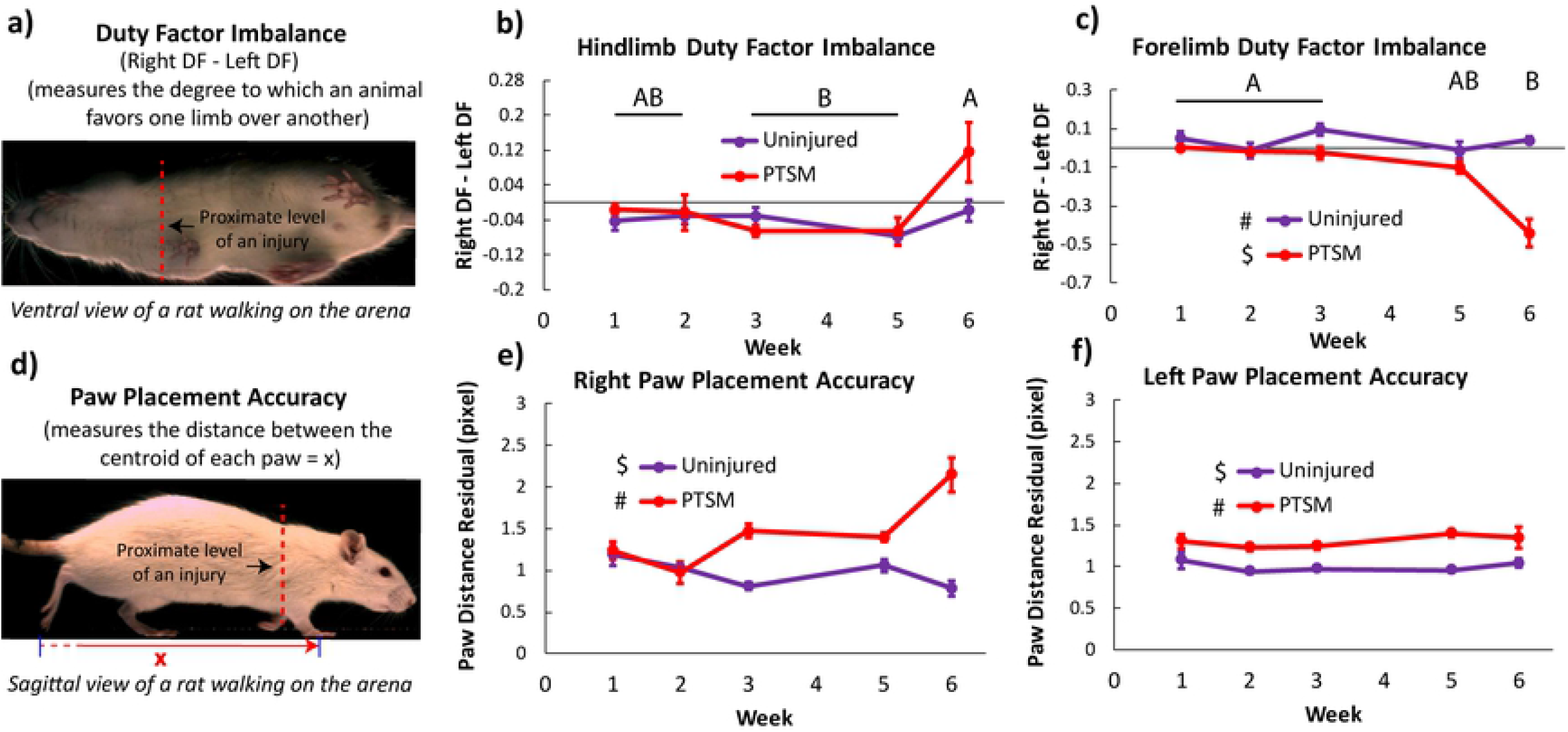
PTSM injury affects the duty factor imbalance of the forelimb and paw placement accuracy of both right and lefts paws. a) A visual definition of duty factor imbalance. b) Hindlimb duty factor imbalance is unaffected by the syrinxes present in the spinal cord due to PTSM injury, whereas c) forelimb duty factor imbalance got affected by the injury model. d) A definition of paw placement accuracy is presented pictorially. Syrinxes present in the spinal cord due to injury greatly affected the right (e) as well as left (f) paw placement accuracy. However, there is no significant difference at the different time points from week 1 to week 6. For all graphs, data are presented as mean ± standard error, for uninjured group n=6, and PTSM group n=9, for both groups subsamples (videos) s=3 to 6. Letters and symbols denote statistical significance as determined by the two-factor ANOVA with Tukey’s *post hoc* (p<0.05).

#### 3.2.1 Duty factor Imbalance and Paw placement accuracy

Duty factor imbalance can be used to indicate the favoring of a limb (either right or left) by a rat while walking in the GAITOR arena. It is evaluated by subtracting the duty factor of the left limb from the duty factor of the right limb as shown in Figure 4a. Duty factor imbalance in hindlimbs was not impacted by PTSM injury as there was no statistical difference between uninjured and PTSM groups (p>0.05), as shown in Figure 4b. However, forelimb duty factor imbalance was significantly impacted by PTSM injury (p<0.05), as shown in Figure 4c, demonstrating PTSM injury caused a detectable locomotion deficit in forelimbs, while not affecting the hindlimbs. The negative duty factor imbalance values indicate that rats spent more time on the left forelimb/hindlimb as compared to the right respective forelimb/hindlimb. A 0 value would indicate no duty factor imbalance such the rat prefers both left and right limbs equally. The forelimb imbalance was amplified from week 1 to week 6 and revealed that animals favored the left forelimb. This suggests a progression of SM over the 6 weeks in the right side of the spinal cord (where the injury was initially made at day 0). Healthy (uninjured) rats should perform ipsilateral fore/hind limb placement similar to every step. To study this pattern we evaluated the paw placement accuracy of both groups using the distance between the centroids of each paw (right forelimb – right hindlimb or left forelimb – left hindlimb) (Figure 4d). Rats with PTSM injury demonstrated reduced accuracy in paw placement for both right and left limbs as shown in Figure 4e,f, indicating the potential interference of the syrinx in observed locomotion deficits. The injury caused the paws to be further apart from each other (positive values of paw placement accuracy which are calculated relative to healthy rats being 1).

#### 3.2.2 Step contact width and step stride length

The width of the step perpendicular to the direction of travel is considered as step contact width (Figure 5a). The step contact width of the hindlimb (Figure 5b) and forelimb (Figure 5c) of PTSM rats was altered as compared to uninjured rats. Interestingly, the forelimb step contact width of injured rats reduced significantly over time, suggesting that PTSM rats were walking cautiously. Moreover, we evaluated stride length, which is the length of a step along the direction of travel, (Figure 5d). Injured rats (PTSM group) exhibited significantly different stride length residual than the rats from the uninjured group in hindlimbs (*p*<0.001) as well as forelimbs (*p*<0.001) as shown in Figure 5 e, f, respectively. The stride length of the PTSM rats was shorter than uninjured rats for both limbs showing an inability of injured rats to place limbs properly on the arena surface, however, stride length was unaffected from week 1 to week 6 (p>0.05). Both parameters, step contact width and stride length of the forelimbs and hindlimbs of injured rats were different than uninjured rats indicating that injured rats were avoiding the full contact of the limbs with the arena surface.

**Figure 5:**
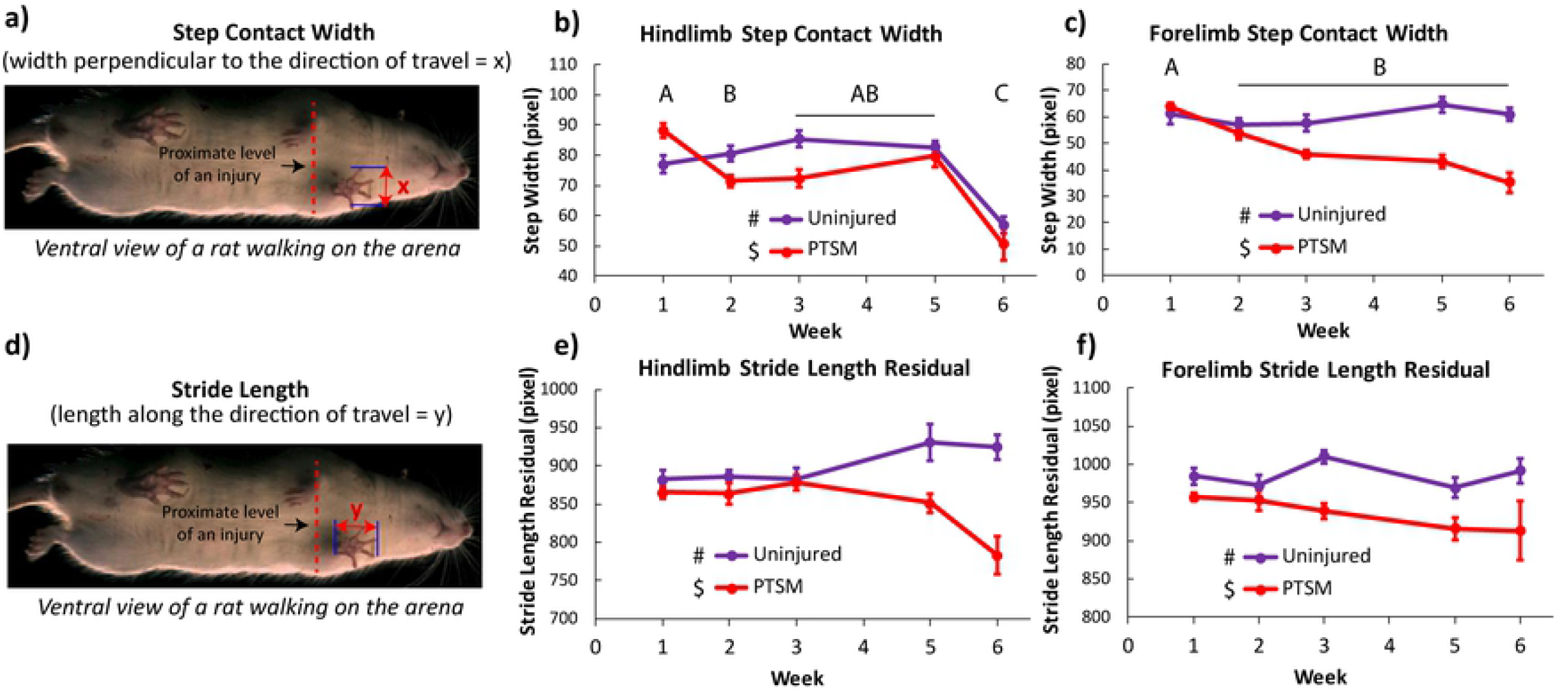
PTSM injury affects step contact width and stride width. a) The definition of step contact width depicted pictorially. b) Hindlimb as well as (c) forelimb step contact width was altered due to PTSM injury. d) Definition of stride length. PTSM injury disturbed the hindlimb (e) and forelimb (f) stride length. For all graphs, data are presented as mean ± standard error, for uninjured group n=6, and PTSM group n=9, for both groups subsamples (videos) s=3 to 6. Letters and symbols denote statistical significance as determined by the two-factor ANOVA with Tukey’s *post hoc* (p<0.05).

### 3.3 Phase Dispersion

Phase dispersion quantifies the degree of coordination during paw placement while rats are walking in the arena. Healthy (uninjured) rats are expected to walk with diagonal paw placement (e.g., right hindlimb – left forelimb and left hindlimb – right forelimb), which corresponds to a phase dispersion of 0. The positive and negative values of the phase dispersion represent that the hind paw is placed after and before the corresponding forepaw, respectively. The right hindlimb – left forelimb and left hindlimb – right forelimb coordination results are presented in Figure 6. The right hindlimb – left forelimb pair shows consistent coordination in uninjured groups, however, PTSM rats lost coordination as indicated by wide phase dispersion after week 1. The PTSM group was significantly different (*p*=0.04) from the uninjured group, with the phase dispersion broadening in week 6 as compared to week 1, further indicating locomotion deficits in PTSM rats over time as shown in Figure 6a. Interestingly, we found no significant difference (*p=0.15*) between uninjured and PTSM groups and over time groups when we compared left hindlimb – right forelimb coordination as shown in Figure 6b.

**Figure 6:**
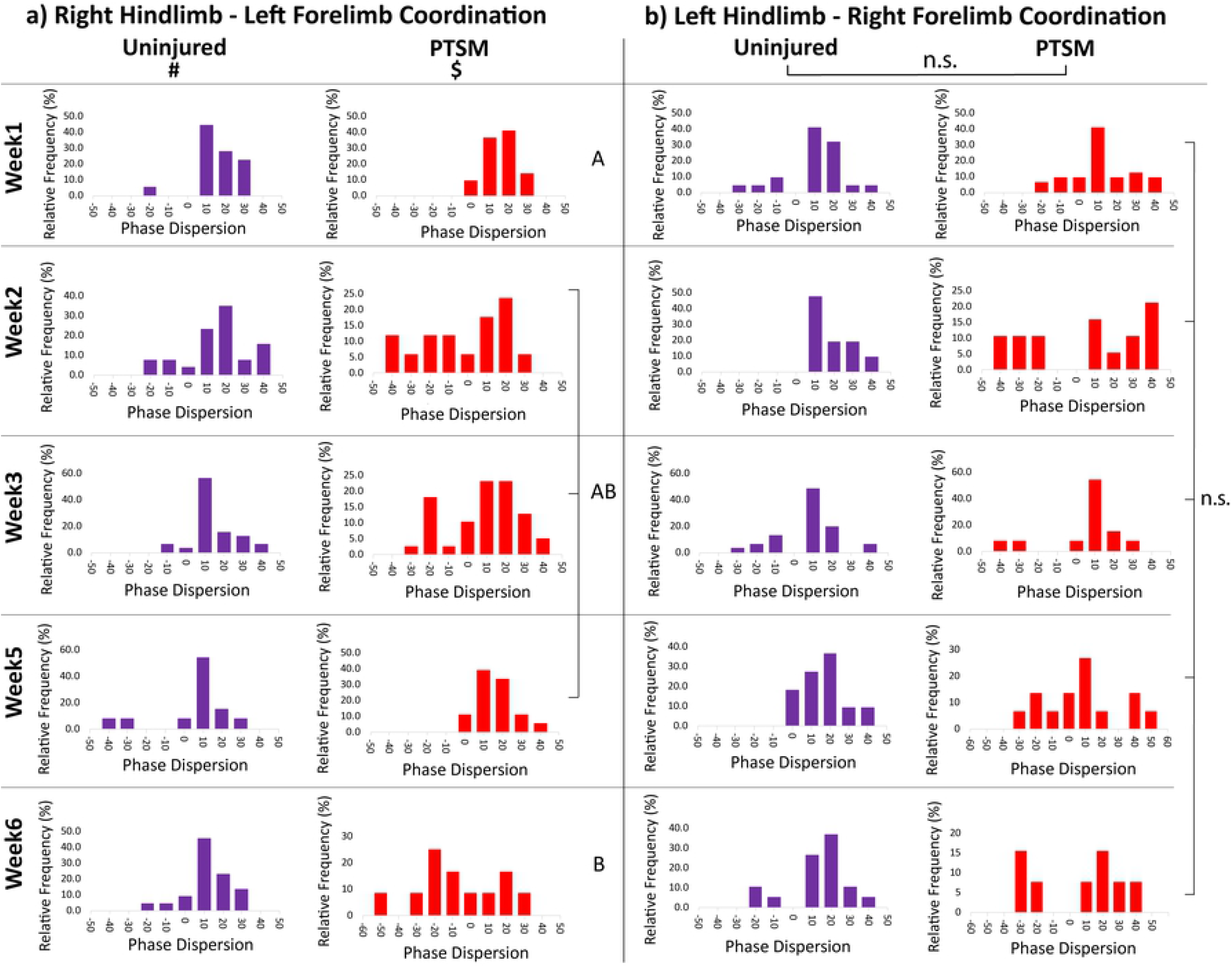
PTSM injury affects diagonal limb coordination of right hindlimb – left forelimb but not left hindlimb – right forelimb. a) The right hindlimb – left forelimb coordination was altered by PTSM injury with wide phase dispersion observed especially at week 6 as compared to week 1. b) There was no effect of PTSM injury on left hindlimb – right forelimb coordination. For all graphs, data are presented as histograms normalized to total step count at that particular time point. For uninjured group n=4-6 and PTSM group n=6-9, and both groups subsamples (steps) s=4 to 6. Letters and symbols denote statistical significance as determined by two-factor ANOVA with Tukey’s *post hoc* (p<0.05).

## 4 Discussion

SM is a devastating neurological disorder that impacts patients’ quality of life. Syrinx progression compresses local neural tissue in the spinal cord, leading to the deterioration of the patient’s condition. To date, only surgical interventions exist to treat SM. Nonsurgical treatments are needed to overcome the risks and challenges of current therapeutic interventions. Unfortunately, there is a poor understanding of the basis for SM, and in response, the molecular aspects of SM have been a recent research focus in the field (12–15). These studies provide a solid foundation for the development of non-surgical treatments for SM; because it is important to establish further analytical tools, particularly those that monitor functional outcomes over time to evaluate treatment efficacy. This work is also an attempt to reveal and quantify subtle functional deficits (especially in locomotion) after PTSM injury in an established rat model. Most SM injury models are not considered to be severe SCI unlike other SCI models such as transection, hemisection, and contusion. Clinically, SM patients show a wide spectrum of symptoms including insidious weakness in the arms and legs, stiffness in the back, shoulders, arms, and legs, loss of sensitivity to pain and temperature over time (6), therefore, in PTSM animal models, there is a need for agreed-upon functional tests to determine and study subtle functional deficits.

In this study, we implemented a modified Hargreaves test, for allodynia and hyperalgesia, to determine the temperature sensitivity of PTSM rats along with uninjured rats as a control (Figure 3). This technique was described first in 1988 and used to determine heat thresholds in the paws of rodents upon application of heat stimulus (26) and further implemented in several animal models (27–31) to quantify heat thresholds corresponding to injury models with or without treatment. The importance and advantages of the method have been highlighted in the literature review (32,33). Unfortunately, our findings showed that the technique was unable to detect allodynia associated with PTSM injury in rats despite being considered as a sensitive allodynia detection technique. Other functional tests such as locomotion outcomes after PTSM injury have been overlooked. Highlighting an outstanding need for reliable, sensitive, and quantitative approaches to study locomotion deficits that likely exist with the PTSM injury in a rat model. To date, existing approaches (such as Basso, Beattie, and Bresnahan (BBB) locomotor rating method) have not reported any such findings. Due to these reasons, we decided to explore GAITOR with AGATHA technique for the rat PTSM model.

Vitally, this study highlights the importance and sensitivity of the recently developed GAITOR with AGATHA technique and its ability to detect subtle locomotion deficits. The relative simplicity of the GAITOR setup and the MATLAB-based open-source AGATHA software is important features that facilitate the implementation of this technique in SCI research (21–23). The ability to observe and capture high-speed resolution images of the paws and to extract meaningful information based on the areas and hues using AGATHA are useful characteristics of the technique that make it sensitive enough to be applied to mild or progressive injury models. This technique also offers the advantage of being an open-source technique offering adaptability, cost-effectiveness, and ease of use over commercially available and closed-source techniques such as DigiGait and Treadscan (22). A main limitation of the GAITOR technique is that animals walking on the area must be walking consistently without changing direction and stopping, or dragging limbs on the arena surface. This makes the technique unsuitable for severe SCI models, which is not an issue for the PTSM rat model. Moreover, this approach requires some level of computer programming knowledge since MATLAB is used to process and extract valuable information from the GAITOR setup’s collected videos. Previously the GAITOR with AGATHA technique was established and successfully implemented to determine locomotion deficit for different animal injury models such as monoiodoacetate injection model of joint pain, a sciatic nerve injury model, an elbow joint contracture model, a hemisection SCI model (21). Also, in a previous study, we effectively implemented this technique to evaluate the effect of intracellular σ-peptide administration on the functional outcomes of rats in a hemisection SCI model (23). Moreover, we extended the use of this technique in a recent study where we found that subcutaneous priming of a novel protein functionalized scaffold notably improved functional outcomes (22). In the last two studies, the importance and the need for a sensitive technique like GAITOR with AGATHA were emphasized showing the ability to obtain more accurate and in-depth quantitative locomotion information, as opposed to a conventional functional test, like BBB, for SCI in rat models.

The results obtained in this study strongly suggest that the AGATHA with GAITOR gait analysis technique is capable of capturing subtle locomotion deficits caused in injured rats due to syrinx formation/expansion in the spinal cord. The duty factor imbalance (Figure 4a) was observed only in forelimbs, suggesting that the location of the syrinx is in the cervical level of the spinal cord (15,25) that impacted the ability of the animal to favor forelimbs equally. The paw placement accuracy results bolster this observation that PTSM injury impacted the left as well as right paw placement accuracy (Figure 4b). This explanation further extends and aligns with step contact width and stride length results as injured rats were avoided proper paw contact with arena surface due to the presence of syrinx in spinal cord tissue as shown in Figure 5a, b. The injured rats demonstrated a lack of diagonal limb coordination in the right hindlimb – left forelimb pair (Figure 6a), moreover, phase dispersion over time became significantly broader in PTSM groups compared to the uninjured group signifies the progressive nature of the syrinx and its impact on locomotion. It is difficult to validate and compare these results from the literature since this is the first study to report locomotion deficit in a PTSM model. However, other functional tests could potentially be used to determine locomotion deficits attributed to PTSM. The forepaw cylinder test (or paw dragging test) (34) is one of the methods that could also be applied since the results from this study show that forepaws were impacted (Figure 4, 5, 6). Additionally, other methods like the rotarod test (35), beam walking test (36), and pole test (37) could also be implemented for PTSM animals in the future to validate the results obtained from the GAITOR with AGATHA technique.

In this paper, we compared the PTSM injured animal group with the uninjured animal group for all locomotion parameters, however, we acknowledge that an additional laminectomy only control is missing to show that the surgical manipulations are not the cause of any observed differences. That being said, laminectomy control animals should not show any differences from uninjured animals if the surgical technique is sound and run by an experienced lab (22).

The results from this study demonstrate the utility of GAITOR with AGATHA for detecting mild and subtle locomotion deficits in a non-severe and indirect PTSM injury small animal model while providing vital quantitative locomotion information. The locomotion parameters presented in this study in terms of duty factor imbalance, paw placement accuracy, step contact width, stride length, and phase dispersion for both forelimbs and hindlimbs, further highlight the sensitivity of the technique for a mild PTSM injury model. The detailed quantitative information provided by GAITOR with AGATHA allowed us to detect subtle locomotion deficits in PTSM injured rats. To the best of our knowledge, this is a first attempt to evaluate this new technique or any technique that is so sensitive, to capture locomotion deficits caused due to PTSM in the rat injury model, which will be useful instrumentation for testing the effects of further treatment development strategies for SM in the near future.

## 5 Acknowledgments

The authors would like to thank Dr. Trevor Ham for training and guidance on GAITOR with AGATHA and help in data processing, as well as Ms. Eleanor Plaster for assistance in video processing. We would like to express our gratitude to Conquer Chiari for providing funding as well as partial funding from Column of Hope.

